# The role of hydrodynamic flow in the self-assembly of dragline spider silk proteins

**DOI:** 10.1101/2022.10.25.513683

**Authors:** Ana M. Herrera-Rodríguez, Anil Kumar Dasanna, Csaba Daday, Eduardo R. Cruz-Chú, Camilo Aponte-Santamaría, Ulrich S. Schwarz, Frauke Gräter

**Affiliations:** Heidelberg Institute for Theoretical Studies, Schloß-Wolfsbrunnenweg 35, 69118 Heidelberg, Germany; BioQuant, Heidelberg University, Im Neuenheimer Feld 267, 69120 Heidelberg, Germany; Institute for Theoretical Physics, Heidelberg University, Philosophenweg 19, 69120 Heidelberg, Germany; Laboratorios de Investigación y Desarrollo, Facultad de Ciencias y Filosofía, Universidad Peruana Cayetano Heredia, Lima, Perú; Interdisciplinary Center for Scientific Computing, Heidelberg University, Im Neuenheimer Feld 205, 69120 Heidelberg, Germany

## Abstract

Hydrodynamic flow in the spider duct induces conformational changes in dragline spider silk proteins (spidroins) and drives their assembly, but the underlying physical mechanisms are still elusive. Here we address this challenging multiscale problem with a complementary strategy of atomistic and coarse-grained Molecular Dynamics (MD) simulations with uniform flow. The conformational changes at the molecular level were analyzed for single tethered spider silk peptides. Uniform flow leads to coiled-to-stretch transitions and pushes alanine residues into *β*-sheet and Poly-Proline II (PPII) conformations. Coarse-grained simulations of the assembly process of multiple semi-flexible block copolymers using multi-particle collision dynamics reveal that the spidroins aggregate faster but into low-order assemblies when they are less extended. At medium-to-large peptide extensions (50%-80%), assembly slows down and becomes reversible with frequent association and dissociation events, while spidroin alignment increases and alanine repeats form ordered regions. Our work highlights the role of flow in guiding silk self-assembly into tough fibers by enhancing alignment and kinetic reversibility, a mechanism likely relevant for other proteins whose function depends on hydrodynamic flow.

## 1 Introduction

Water plays a major role for the stabilization, structure and dynamics of proteins [1, 2], mainly via hydrogen bond networking [3] and screening of electrostatic interactions [4]. Although most proteins function in the context of a quiescent fluid, there are important situation in which hydrodynamic flow impact protein structure by inducing non-uniform drag force along the protein. Hydrodynamic flow can mediate a large variety of processes such as material synthesis, blood coagulation and protein misfolding [5, 6, 7]. For example, the flow-induced activation of von Willebrand factor (vWf) protein is crucial in hemostasis [8, 9, 10, 11]. Shear and elongational flows can lead to protein unfolding, misfolding and aggregation, and can accelerate fibrillation of amyloid-β peptide [12, 13, 14, 15]. A complete molecular understanding of how proteins respond to flow, however, remains elusive.

Here we study an important example from material synthesis, namely assembly of dragline spider silk proteins (spidroins) into a fiber, which occurs in the *Major Ampullate* (MA) spinning gland and outperforms mechanically any biomaterial made by nature [7, 16, 17, 18, 19, 20, 21, 22]. Major ampullate spidroins consists of pH-dependent folded N-terminal and C-terminal domains, which experience conformational changes from pH 7.0 down to pH 5.0 in the spider spinning gland, thereby initiating assembly [20]. Inbetween these terminal domains, spidroins feature a long repetitive region, which makes up roughly 90% of their total sequence (Figure 1a) [23]. This repetitive region of silk proteins comprises alternating blocks of hydrophobic alanines, which form *β*-sheet stacks in the silk fiber, and hydrophilic glycine-rich repeats, which form the amorphous and extensible matrix of silk [16, 22, 24]. A large body of work explored the dependence of silk fiber assembly under elongational and shear flows using microfluidics [6, 18, 25, 19, 20, 26]. In single molecule force spectroscopy experiments, single spidroins molecules largely followed a worm-like chain behavior, but also exhibited unique unfolding steps [24, 27, 28, 29]. However, flow-dependent dynamics of single silk proteins and flow-induced spidroins self-assembly remain largely unknown.

**Figure 1:**
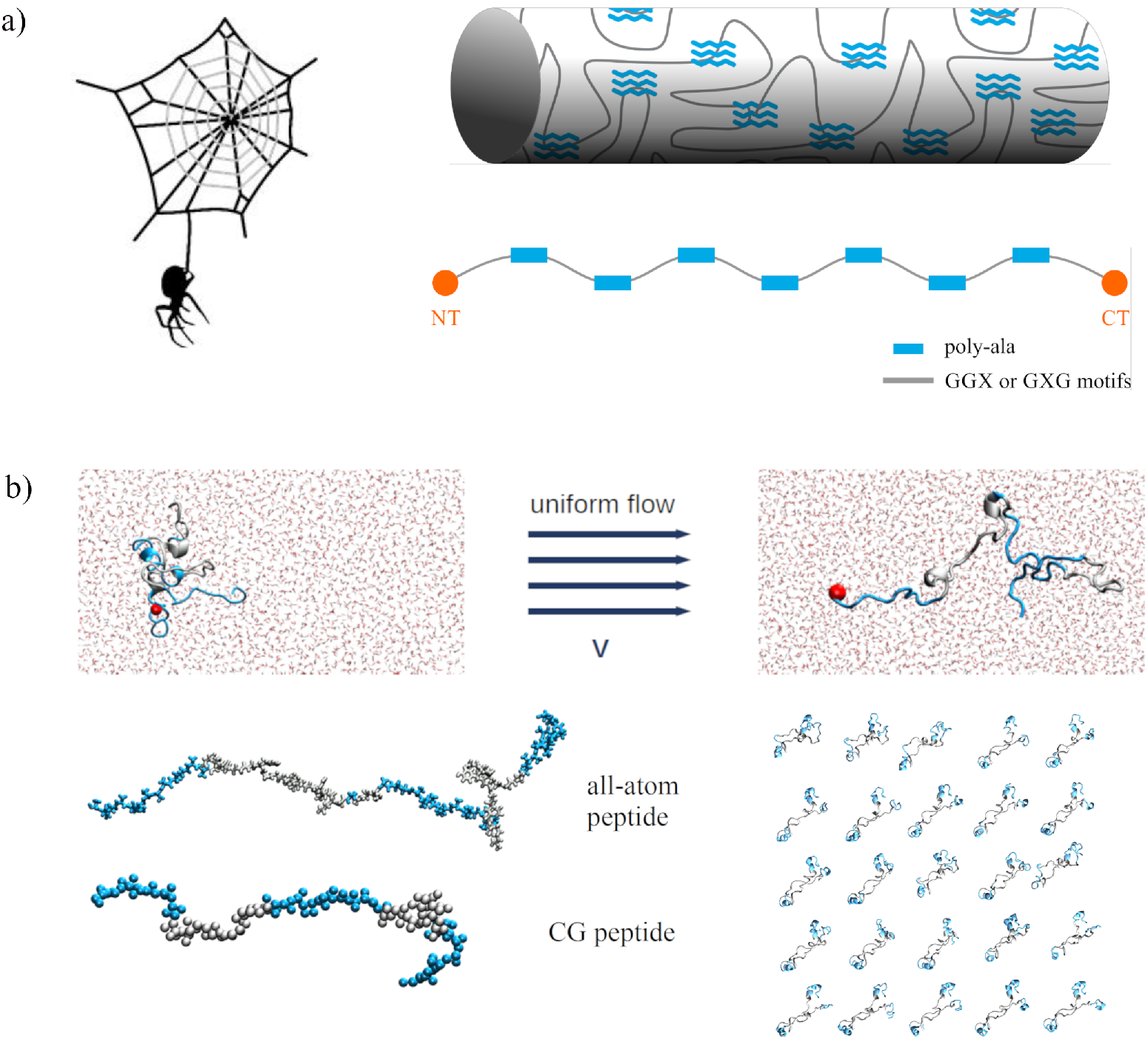
Dragline silk and computational model for spidroins under uniform flow. (a) Dragline silk corresponds to the structural part of spider’s webframe (top left), which fiber microstructure consists of embedded *β*-sheet crystals into an amorphous viscoelastic matrix. The fiber is formed upon directed SA induced by flow of the spidroins, an scheme of the *MA Euphrosthenops Australis dragline spidroin* sequence is shown. Its molecular structure comprises three regions: pH dependent N- and C-terminal domains (NT and CT, red), as well as a long repetitive region with alternating poly-alanine (Poly-Ala, blue) repeats and glycine-rich motifs (GGX or GXG, with X being another amino acid, grey). (b) The MD model of silk under flow considers atomistic (all-atom) and coarse-grained (coarse-grained) models of a fragment of the repetitive region, containing three poly-Ala and two amorphous regions, resulting in ~ 80 amino acids in total. Spheres represent atoms (for all-atom) or backbone and sidechain beads for each amino acid (coarse-grained). The spidroin fragment is tethered at the N-terminus (red sphere), and a uniform flow is produced to stretch the peptide. To model SA under uniform flow, sets of 25 of peptides (bottom right) are tethered along the flow direction and able to move transversely. Color coding as in the spidroin scheme in (a).

Unraveling the physical effects of elongational flow during formation of spider silk would fill an important gap in our understanding of the assembly pathway of the spidroins. In particular, it could help to solve the open question whether the spinnability of silk requires intermediate formation of micelles or not [30, 31]. Additionally, it could help to better describe liquid crystalline flows inside the s-duct [17, 32, 33]. The lack of a more detailed understanding of silk self-assembly is, among others, due to the challenge to experimentally work with the highly aggregation-prone silk proteins. In principle, the proteins should be soluble to prevent fatal aggregation. It is difficult to experimentally manipulate the silk dopes due to their high concentrations ~ 50%wt/v and their sensitivity to changes in the micro-environment (pH, ion conditions and shear forces). Moreover, the lack of structuring of the silk monomers in most of its domains limits the use of structural studies.

Theory and simulations have shed further light into the stretching and assembly behavior of biopolymers under flow. For the simplest case of biopolymers under uniform flow, Monte Carlo simulations of tethered DNA chains revealed that the nonlinear elastic dumbbell model predicts their deformation [34]. The simulations included Brownian motion, entropic elasticity, and variations in the drag coefficient while the chain is deformed. Brownian dynamics simulations of DNA molecules in steady flow showed interesting differences between WLC and Rouse models in a strong steady flow, with convective as opposed to diffusive propagation of tension along the chain, respectively, leading to a distinct time evolution of chain extension [35]. In addition, bead-spring models have proven useful for describing (bio)polymers, including proteins under flow conditions. A prominent example is the coarse-grained (CG) modelling of vWf multimers in shear flow, with one bead representing roughly one monomer and a Lennard-Jones potential to account for average attractive interactions [36, 37]. A variety of methods have been used in such mesoscopic simulations for the hydrodynamic interactions of the polymer with the solvent, based on Brownian Dynamics (BD) [9, 11, 36, 38], multi-particle collision dynamics (MPCD) [39, 40, 41, 42], dissipative particle dynamics (DPD) [19, 37, 43], or lattice-Boltzmann [44]. In the context of spider silk SA by flow, a mesoscopic modelling using DPD served to find that intermediate ratios of hydrophobic and hydrophilic blocks observed in spider silks lead to exceptional silk fiber formation [19]. Shear flow was shown to lead to enhancement of β-sheet crystals, confirming a previous experimental research[6, 7, 18]. However, the effect of hydrodynamic flow on the structure of silk fibers remains elusive.

In contrast to these mesoscale simulations, atomistic MD simulations can give detailed insight into the conformational changes of proteins induced by flow, such as ubiquitin unfolding [45] or secondary structure transitions in the platelet receptor glycoprotein Ib [46]. The high degree of detail offered by atomistic MD in explicit solvent comes at a high computational cost, resulting in high flow rates used in these studies. Additionally, the high computational costs also limits the possibility to study protein assembly processes like spider silk protein self-assembly, thus most of the studies are associated with single protein dynamics under flow.

In this study, we attack these open questions in a two-pronged approach of simultaneously performing atomistic and coarse-grained simulations at physiologically relevant flow velocities. As representative example, we chose an ~80 amino acid-long sequence from the repetitive part of MA *Euphrosthenops Australis* dragline spidroin, which consists of three poly-ALA repeats, and two amorphous regions (Figure 1a). We modeled these proteins at the mesoscopic scale as a simple block-copolymer using a Go-like potential [47] combined with Multiparticle-Collision Dynamics (MPCD) [39] to model a uniform flow. For comparison, we performed atomistic MD simulations of the same spidroin fragments in uniform flow of explicit water [48]. We find the proteins to largely follow the expected freely-jointed chain behavior as known from single molecule stretching experiments and simulations, though with interesting differences. Flow unravels long-range poly-alanine interactions and promotes *β*-sheet and poly-proline II conformations, both known to be present in silk fibers, in particular within the poly-alanine repeats. Regarding self-assembly, we found an enhancement of oligomerization at low flow rates. The alanines are more prone to form interchain contacts than the residues in the amorphous region and this favors formation of *β*-sheet crystals during fibrillation. Our results give molecular insight into the drag force and stretching dynamics of disordered and unfolded proteins in flow as well as into flow-induced assembly of spider silk and related protein-based materials.

## 2 Methods

In all simulations, we model a representative fragment of silk spidroin, consisting of three polyalanine repeats separated by two glycine rich regions, with a total length of ~80 amino acids. Figure 1b shows all-atom and CG representations of the repetitive fragment we simulated. The simulations comprise both single silk peptides, at both all-atom and coarse-grained level, and multiple silk peptides, only at coarse-grained level, under uniform flow.

### All-atom MD simulations of single peptides

We used the Gromacs 2018 version [49]. The sequence of the peptide originates from the repetitive part of the *Euprosthenops Australis* spidroin at the MA spinning gland and is: [A]_13_GQGGQGGYGGLGQGGYGQGAGSS[A]_14_GRGQGGYGQGSGGN[A]_12_. It is organized in alternating Poly-Ala repeats and glycine-rich disordered fragments. We used the Amber force field Amberff99sb-star-ILDN in combination with the Tip4pD water model, which previously proved appropriate to model intrinsically disordered proteins such as the one investigated here [50, 51, 52]. Periodic boundary conditions were used in all three dimensions for all simulations. The concatenated simulation time including system preparation and production runs with flow is 18.5 μs. Before introducing uniform flow, we prepared and equilibrated the system as follows: an energy minimization with 100000 steps starting from a randomly chosen conformation of the protein was followed by 5 ns of MD simulations in the NVT ensemble and 10 ns of MD in the NpT ensemble with the protein with position restraint. Temperature coupling was done by a V-rescale thermostat during equilibration and production runs [53]. The Parrinello-Raman barostat [54] was used for pressure coupling in the NpT runs. LINCS constraints were used for all bonds to allow a 2 fs integration step. After preparation, we ran 300 ns in the NpT ensemble to equilibrate the protein without restraining the protein. After protein equilibration, we took 10 different conformations and prepared larger simulation systems along the flow direction. We simulated 10 flows with mean velocities from 0.0 m/s up to 0.48 m/s, each for 600 ns in an NVT ensemble. During the simulation we pulled waters inside a slice of 4 nm size with a constant force, using a modified Gromacs version to introduce uniform flow [48]. The V-rescale thermostat was used, which relaxed the system to achieve a constant velocity flow given by the externally applied velocity. The peptide was placed 4 nm away from the pulling region to avoid undesired stretching not associated with the flow, and artifacts due to the pressure drop inside the slice and at its interface. For the production run, the first alpha carbon of the peptide was position restraint to prevent translation. A timestep of 2fs and LINCS constraint on h-bonds were used. We simulated for three different replicates for every flow velocity, each starting from a different velocity seed.

### Coarse-grained spidroin model

We simulated the same system described in the previous section at coarse-grained level, i.e. at the resolution of two beads per amino acid (backbone bead and sidechain bead) rather than atoms. The two beads are located on the *C_α_* and *C_β_* atoms of each amino acid. The peptide is simplified as a block copolymer composed of two types of amino acids: alanine and a non-specific amino acid type for the disordered region. The amount of Ala residues in every region is the same as in all-atom simulations. Since the primary intermolecular interaction within silk fibers are *β*-sheets formed from the Poly-Ala regions, we considered attractive (native) interactions only between Ala residues in our model, more specifically between pairs of alanine backbone beads (representing the backbone hydrogen bonding within *β*-sheets) and between pairs of alanine sidechain beads (representing hydrophobic packing between methyl groups across *β*-sheets [55]. These ‘native’ interactions were modelled as attractive Lennard Jones interactions and all other bead-bead interactions as purely repulsive terms. We now describe all interactions of our spidroin mesoscopic SOP model including sidechains (SOP-SC) in detail.

The total potential energy *U*_total_ to describe the interaction between these monomers is based on the SOP-SC polymer model [47], is defined as

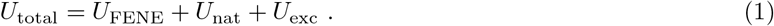

The first term *U*_FENE_ is a modified form of Finitely Extensible Nonlinear Elastic (FENE) potential for the bonded interactions, i.e. beads are connected by nonlinear springs. The summation runs over all bonded backbone-backbone (BB) and backbone-sidechain (BS) beads. The potential is:

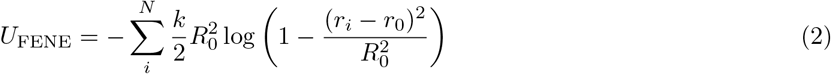

where *r_0_* is equilibrium length of the bond, *R_0_* is the maximum extent of the bond and *k* is the spring constant. The second term in equation (1) represents native interactions between the alanine pairs *i,j* such that |*i* – *j*| > 2. It is modeled with Lennard Jones (LJ) potential with equilibrium distance *σ* and energy *ϵ*:

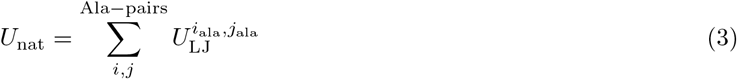

with

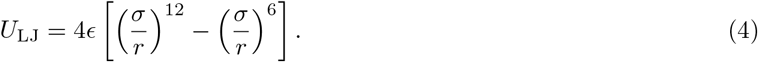

The distance between BB and sidechain-sidechain (SS) beads is 0.5 nm, corresponding to the distance when there is *β*-sheet secondary structure formation in silk. Therefore, *σ* = 0.5 nm. BS interactions within alanines are not treated as attractive interactions, but are included in the repulsive term (see below).

The last term represents exclusion interactions:

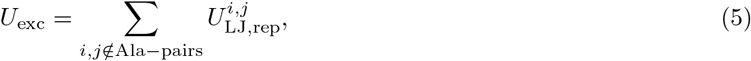

and

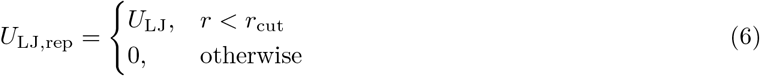

where *r*_cut_ = *r_m_* is the distance at which the potential has a minimum value. The summation in equation 5 runs over all possible pairs of beads, except over native interaction pairs.

The parameters associated with the potential energy are for:

- *U_FENE_* potential: bond length *r*_0_ = 1.0 [*a*], *k* = 500 [*k_B_T*/*a*^2^] and *R*_0_ = 0.5 [*a*].
- *U*_nat_ potential: *ϵ_nat_* = 0.25 [*k_B_T*] and *σ_nat_* = 2 *r*_0_ [*a*] of the LJ potential for both BB and SS interactions.
- *U*_exc_ potential: the parameters have the same value for BB, SS and BS interactions, which are *ϵ_exc_* = 1.0 [*k_B_T*] and *σ_exc_* = *r*_0_ [*a*].

### Multi-particle Collision Dynamics simulations

The CG model of the peptide was combined with hydrodynamic flow using Multiparticle Collision Dynamics (MPCD), which is a particle-based mesoscopic simulation technique for the Navier-Stokes equation. It incorporates hydrodynamic interactions and thermal fluctuations [56, 39]. We employ a local cell-level scaling thermostat [57] to maintain the temperature of the system constant. The fluid in MPCD consists of point particles *m*, position 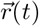 and velocity 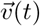. Particles are sorted into a cubic lattice with a lattice constant *a* and subjected to the dynamics of two steps:

- a streaming step where the particles move ballistically during a collision time step Δ*t_c_*. Particle positions at every timestep are described by

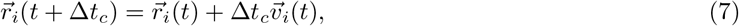

where 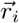 and 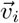 are the positions and velocities of every particle. Position update follows thus a standard integration of Newton’s equation of motion
- The collision step where particle velocities are altered. The center-of-mass velocity of the cells 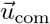 is computed, and subsequently the relative velocities of the particles with respect to 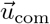 are rotated around a random axis by an angle *α*, which ensures linear momentum conservation and thermal fluctuations. The velocity evolution for every particle in time is given by

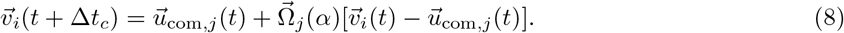

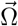 and *j* correspond to the rotation matrix in spherical coordinates and the cell index respectively. The beads of the peptide are included in the MPC collision step, in order to take into account solvent-peptide interactions. The integration time step Δ*t* was chosen to be smaller than the collision time step Δ*t_c_*. To induce flow, a constant force *F* was applied to every solvent particle.

The fluid viscosity *η* in MPCD depends on the collision time step Δ*t_c_*, particle density *ρ* and mass *m*. It has two contributions: collision viscosity *η_col_* which results of momentum transfer between the particles during the collision step and kinetic viscosity *η_kin_* which results from the stage of particle streaming, and are given by

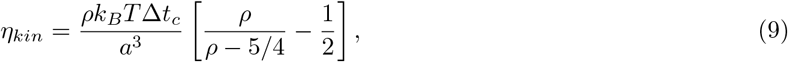

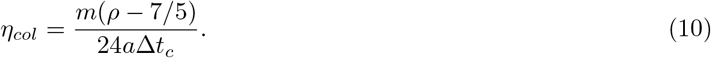

For low Reynolds numbers *η_kin_* can be neglected and the main contribution to the total viscosity *η* is provided by the collision viscosity *η ≃ η_col_*

The simulation parameters are expressed in basic units *k*_B_*T* = 1, *a* = 1 and *m* = 1, where *k*_B_ is the Boltzmann constant and *T* is the temperature.The box volume is *V* = 100.0 × 50.0 × 50.0 *a*^3^ and the particle density is set to *ρ* = 10. The mass of every monomer is *M* = *ρ*m, the integration time step Δ*t* is 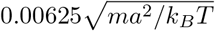 and the collision time step is 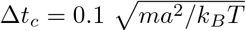.

The simulations consisted of an equilibration of the system for 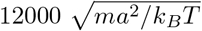 starting from a fully extended chain, from which its final conformation was taken to simulate under 10 different flows. The force applied to the solvent particles to simulate uniform flow at a range of velocities was in the interval of (5 ·10^−07^ – 2.7 · 10^−4^)[(*k_B_T*/*a*)]. We carried out three replicas with different velocity seeds for each flow velocity in both single peptide and multiple peptide simulations.

### Coarse-grained simulations of silk assembly

To monitor the self-assembly of spider silk proteins, we performed simulations of peptides sets under uniform flow in coarse-grained simulations. Every spidroin corresponds to a random conformation taken from the single peptide simulations in a given flow regime. Like in the single silk spidroins simulations, the silk peptides are tethered as well along the flow direction. Initially, the peptides are placed equi-distant from each other in a matrix of 5×5, where the positions (*y, z*) of the tether atoms correspond to the nodes of the matrix. Figure 1b (bottom right) shows the initial configuration of the peptides for one replicate of the all-atom simulations. The peptides are allowed to move transversely (in the *yz* plane) and are tethered at the same position in the flow component *x*.

To monitor the self-assembly of spider silk proteins, we performed simulations of peptides sets under uniform flow in coarse-grained simulations. Every spidroin corresponds to a random conformation taken from the single peptide simulations in a given flow regime. Like in the single silk spidroins simulations, the silk peptides are tethered as well along the flow direction. Initially, the peptides are placed equi-distant from each other in a matrix of 5×5, where the positions (*y, z*) of the tether atoms correspond to the nodes of the matrix. Figure 1b (bottom right) shows the initial configuration of the peptides. The peptides are allowed to move transversely (in the *yz* plane) and are tethered at the same position in the flow component *x*.

| Mean flow velocity<br>mpcd units | Mean fractional extension of peptides<br>L | Simulation time<br>mpcd units | Number of replicates | Number of peptides |
| --- | --- | --- | --- | --- |
| 0.1 | 0.4 | 800 | 3 | 25 |
| 0.2 | 0.7 | 700 | 3 | 25 |
| 0.3 | 0.8 | 600 | 3 | 25 |
| 0.4 | 0.9 | 600 | 3 | 25 |
| 0.5 | 0.4 | 500 | 3 | 25 |
| 0.7 | 0.7 | 300 | 3 | 25 |
| 0.8 | 0.8 | 200 | 3 | 25 |
| 1.0 | 0.9 | 200 | 3 | 25 |

We used the same protocol and simulation parameters as in the single peptide case (Subsection 2) except the bond length was decreased to *r*_0_ = 0.7 from *r*_0_ = 1.0 to reduce the overall system size and increase computational speed. This only alters the effective friction of a single bead. The initial peptide separation is 7,6 nm and the system concentration is ~ 4%wt/v for the CG system. In total, we performed 6 simulations with 3 replicates each. A description of each simulation is contained in Table 2.

### Persistence and Kuhn lengths

To obtain the Kuhn length of the polymer at all-atom and coarse-grained scale, we fitted an exponential function to the auto-correlation function *C*(*s*) of the silk peptide backbone segments s, using:

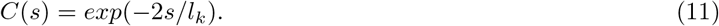

The Kuhn length for all-atom and coarse-grained simulations was 0.2 nm and 1.6 *a* respectively. With *l_k_* = 2*l_p_* we obtained the Kuhn length for both models.

### Drag force calculation

We calculated the drag force acting on the protein in all-atom MD simulations from inter-atomic forces *F_ij_* between protein atom *i* and protein atom *j*, summed the *x*-component of all these forces up (with *x* being the flow direction), and averaged over time. We used the same definition of the drag force for the coarse-grained simulations, just that in this case, the forces only included the momenta transferred in the collision steps as only local protein-solvent forces are considered in MPCD.

### Secondary structure analysis

For the secondary structure analysis, we defined residues to be in a *β*-sheet conformation when the dihedrals fall into the intervals of 90 ≤ *ψ* ≤ 180 and −180 ≤ *ϕ* ≤ –90 (circles), and in a PPII conformation if 128 ≤ *ψ* ≤ 180 and –90 ≤ *ϕ* ≤ –58 (squares). In order to observe the relevance of alanines in the formation of *β*-sheets we show the percentage of Ala and non-Ala residues like: %*Ala_β/PPII_* = *N_Ala_β/PPII__* * 100/*N_Ala_* (blue symbols) and %*nonAla_β/PPII_* = *N_nonAla_β/PPII__* * 100/*N_nonAla_* (gray symbols). *N_Ala_β/PPII__* and *N_Ala_* correspond to the number of Ala residues in *β*-sheet or PPII helix and the total number of alanines in the peptide sequence, while *N_nonAla_β/PPII__* and *N_nonAla_* are the number of residues of the amorphous region in *β*-sheet or PPII and the total number of residues different from alanines.

### Friend of friends algorithm

To monitor whether a silk peptide forms part of an oligomer of certain size or not, we adapted a group selection algorithm called Friend of Friends (FoF) to our case [58]. First, we chose a peptide, which has not been previously assigned to an oligomer group *i*. Next, we search for partners of this peptide based on the inter-chain *C_α_* carbons contact formation, where the condition to have a contact of two atoms between peptide 1 and peptide 2 is defined as 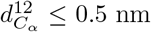 (see the spidroin inter-contact sketch at Figure S1). In case, the peptide does not have a partner, we assign it to an “isolated” peptide list. When partners are found, all of them are added to the list of *i* oligomers. We search repeatedly until no further members can be found. This process runs over all frames of the simulation. Therefore, we can monitor not only the time evolution of individual chains that form oligomers, but also their contact formation with other chains.

## 3 Results

### 3.1 Single silk spidroins under flow

We first asked how the flow velocities are related to the drag force acting on the peptide. As the drag force changes dynamically with the peptide conformation, a simple analytical model of the drag as a function of velocity might not be appropriate. We instead calculated the drag force directly from the forces between the solvent particles and the protein particles (atoms or beads respectively). In the all-atom case, we computed the vector-sum of the non-bonded forces between all atoms of the water molecules and of the peptide over time. In the coarse-grained case, we obtained the drag force from averaging the collisions of the solvent particles with the beads, as the solvent-protein interactions solely occur through local momentum exchange. In both all-atom and coarse-grained simulations, the force components along the flow yields a non-zero average force, the drag force. We averaged the drag force over the last half of each trajectory and over all three replicates. Interestingly, independent on the resolution of the simulations, i.e. for both all-atom and coarse-grained, we obtain an overall linear increase in drag force with flow velocity (Figure 2a and Figure 2b). Thus, the drag force effectively follows Stokes’ law, with an effective hydrodynamic radius of 2.52nm (obtained from the linear fit to the AA data of Figure 2a). We conclude that our spidroin peptide does not exhibit a pronounced globule stretch transition at a critical flow speed, which would lead to a deviation from a linear drag-flow relationship, but instead steadily extends with increasing flow velocity. This is in sharp contrast to the behavior seen previously for proteins in shear flow, which showed a rather abrupt globule-stretch transition (in case of VWF) [9] or a one-step unfolding (in case of ubiquitin) [40, 41], respectively. However, a gradual stretching through multiple unfolding intermediates were observed for ubiquitin in the case of elongational flow, [40, 41] directly in line with the gradual extension of the spidroin with increasing flow velocities.

**Figure 2:**
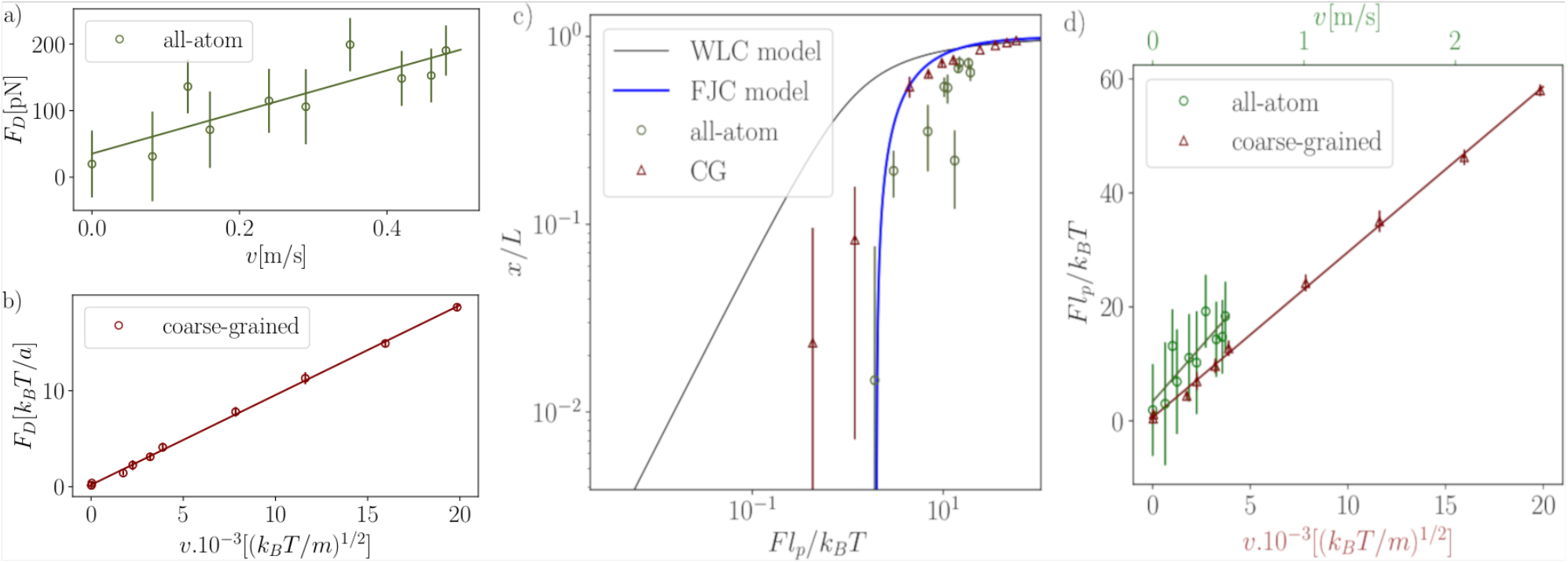
Stretching silk peptides under uniform flow. (a) Averaged peptide drag force over mean flow velocity for all-atom simulations. The drag force in the peptide originates from the non-bonded interactions between the water and protein atoms. The green solid curve is a linear fit. (b) Averaged peptide drag force for coarse-grained simulations, the drag force was computed from the collisions between the solvent particles and the amino acids of the peptide. The maroon solid curve is a linear fit. (c) Dimensionless peptide drag force as a function of the normalized peptide extension, *L* is the contour length. Circles and triangles correspond to all-atom and coarse-grained simulations, respectively. The persistence length was extracted from fitting to equation (11). Black and blue solid curves: normalized theoretical prediction from the WLC and FJC models, *f_FJC_*(*x*) = 1/(1 –*x*) and *f_WLC_*(*x*) = 1/4(1 –*x/L*)^-2^–1/4 +*x/L*. Error bars in all the plots are standard errors of the mean for nine data points, obtained from time-averages over three 100 ns windows of the second half of each of the three trajectories per flow speed. (d) Dimensionless drag force over flow velocity for all-atom and coarse-grained simulations, *lp* is the persistence length. The highest velocity of the atomistic simulations is about one fifth the highest velocity of the coarse-grained simulations. The maroon and green solid curves are linear fits.

To quantitatively compare the all-atom and coarse-grained scales, Figure 2c shows for both sets of simulations the dimensionless drag force in the peptide as a function of the normalized extension along the flow. *L* and *lp* correspond to the contour length and persistence length, which are 24.3nm and 0.4nm for all-atom simulations as well as 80 *a* and 3.1 *a* for coarse-grained simulations. We recover a non-linear force-extension curve typical for polymers when stretched by a pulling force. The all-atom model yields overall lower extensions for a given drag force, but given that they are fully independently parameterized, the agreement is satisfying. The coarsegrained model extends more readily is likely due to our approximation that only attractive interactions are present within alanine residues, neglecting unspecific favorable interactions involving non-alanine residues of the disordered region. The force-extension behavior is reminiscent of the behavior predicted by the freely-jointed chain (FJC) [59] or worm-like chain (WLC) model [60] of polymers. Indeed, we find a good agreement of our data with the FJC model (solid blue curve Figure 2c). We find slightly less agreement between the WLC model and our particle-based simulations (Figure 2c, solid black line), which we explain by the nature of the protein backbone the (albeit only partially) rotatable bonds of which are better depicted by a FJC model than by the rod-like WLC with uniform bending stiffness. Stretching the silk peptide to ~80 % of its contour length (*x/L* = 0.8) requires a drag force from uniform flow in the range of 100-200 pN, according to both all-atom and coarse-grained simulations. Interestingly, this range covers the rupture forces of ~176±73pN required to stretch single dragline silk molecules by an atomic force microscope [27]. To estimate the velocities achieved in the coarse-grained model, the Figure 2d shows the normalized drag force as a function of velocity for all-atom and coarse-grained models, the velocities in the all-atom model are re-scaled by a factor *α* = 7.75. The highest flow velocity in the coarse-grained simulations corresponds to ~2.6m/s.

Alanine residues are expected to play an important role during shear-induced unfolding of silk peptides and their assembly into a fiber. To analyze the relevance of Ala residues in the silk protein conformational ensembles in flow, we quantified the number of Ala contacts along flow velocities in all-atom (Figure 3a) and coarse-grained simulations (Figure 3b). In the all-atom case, we observe a drop of Ala-Ala contacts between 0.16 and 0.24 m/s of flow rate. Also the fractional extension of the all-atom peptide as a function of the flow rate confirms this behavior (Supplementary Figure S2a): initially the peptide is extended up to about 30% of the contour length from 0.08 m/s up to 0.16 m/s. After this threshold, the peptide is stretched 20% further in the mean flow velocity interval of (0.24 - 0.29) m/s up to around 80% of its contour length at 0.48 m/s.

**Figure 3:**
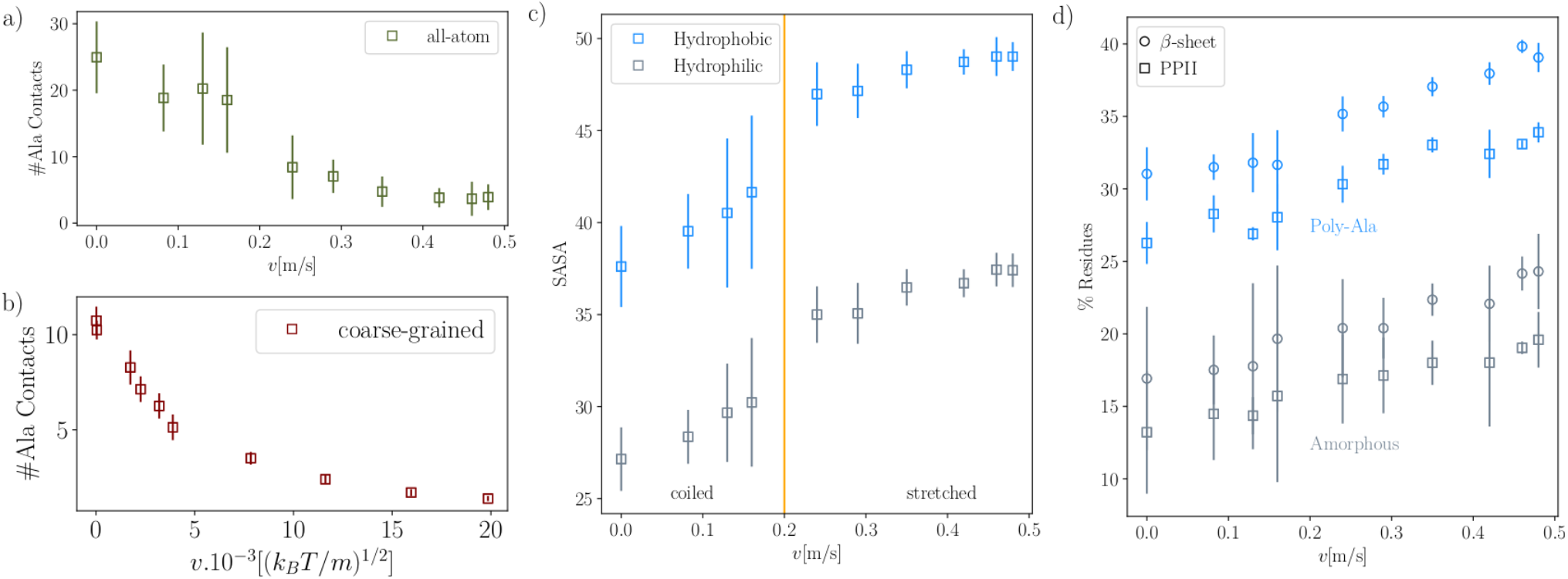
Molecular details during silk elongation. a) Number of contacts of Ala residues as a function of time in all-atom MD simulations. The cutoff radius within we examine a contact formation is 0.5 nm. b) Number of contacts of Ala residues as a function of time in coarse-grained MD simulations. The cutoff radius within we examine a contact formation is 0.5 nm. c) Hydrophobic and hydrophilic solvent accessible surface areas over flow rate for all-atom simulations. At about 0.2 m/s a transition of the peptide to a extended state is observed (solid vertical orange line). The highest velocity of the atomistic simulations is about one fifth the highest velocity of the coarse-grained simulations. d) Flow populates *β*-sheet and PPII conformations. Percentage of dihedral angles of amino acids in *β*-sheet (circles) and PPII conformation (squares) as a function of mean flow velocity, as observed in all-atom simulations. The data is separated into alanine (blue symbols) and amorphous regions (gray symbols), and normalized by the total number of residues in these two regions. Error bars show the standard error of the mean. See Methods for the definition of the dihedral intervals for both secondary structures.

In the case of the coarse-grained peptide we observe a less abrupt transition to a fully extended state without Ala-Ala contacts. The contacts steadily decrease from 10 contacts to nearly 0 contacts between 0.02 and 20×10^-3^(*k_B_T*/*m*)^1/2^(Figure 3b). The number of total and amorphous contacts over mean flow rate in Figure 3b shows the same tendency. With regard to the coarse-grained fractional extension (Supplementary Figure S2b) over the mean velocity, Ala contacts are maximal only at 0-10% fractional extensions, and are largely lost only beyond 90% extensions. Thus, the CG model does not reproduce the comparably sudden transition observed at all-atom. In addition, the coarse-grained model shows less contacts initially, i.e. at low flow velocities (~12 instead of ~24 for all-atom). We conclude that the CG model reproduces the overall loss of contacts in flow, though it cannot resolve the collapse-stretch transition of all-atom simulations.

The observed abrupt transition of the peptide from coiled to stretched conformation in the all-atom is associated with a decrease of the hydrophobic effect. It can be observed at the Figure 3c, which shows the solvent accessible surface area (SASA) over flow rate. Upon force, the hydrophobic residues are not buried and the silk peptide suddenly transits from a coiled to a extended state. This transition occurs about at 0.2 m/s of flow rate (see the orange vertical line at Figure 3c).

The formation of *β*-sheets is a key factor in the self-assembly of spider silk and requires flow [18]. However, how flow induces *β*-sheet formation is poorly understood. In recent studies, a significant population, namely ~ 24%, of poly-proline II (PPII) helix in the repetitive region of MA spidroin dopes, i.e. prior to flow-induced elongation during spinning, from the *Euphrostenops Australis* spider has been observed [61]. It was suggested that PPII helices can form a rigid structure that can be quickly transformed into *β*-sheets. We here asked to what extent our spidroin fragment samples *β*-sheet and PPII conformations and how these backbone propensities are influenced by flow. We defined for each amino acid the respective backbone configurations based on its two backbone dihedral angles (see Methods for details). We averaged over all alanines and all amorphous-phase residues (non-alanines) and normalized by the total number of alanines and non-alanines, respectively. We note that we analyzed the secondary structure propensities only for the AA simulations, as the CG lacks the required resolution.

Figure Figure 3d shows the percentage of amino acids with dihedral angles either in *β*-sheet or PPII conformation as a function of mean flow velocity. The percentage of alanines in both, *β*-sheet and PPII conformations is increasing with the mean flow velocity for both the poly-Ala and amorphous regions. Both, *β*-sheet and PPII conformations populate the upper left corner of the Ramachandran plot and are close to the most extended backbone configuration with *ψ* = 180 and *ϕ* = –180. For this reason, flow is expected to drive the amino acids into this region of the dihedral space, as previously also observed for disordered proteins under a stretching force [62]. The content of the backbone torsion angles in *β*-sheet and PPII in the poly-Ala regions remains around 10% higher than in the amorphous regions, largely independent of the flow velocity. This higher ratios are in line with the view that the poly-alanine repeats preferentially drive *β*-sheet formation, i.e. crystallization, during fiber formation.

In both the alanine and amorphous phase, *β*-sheet conformations are slightly preferred over the PPII conformation, and they together represent the majority of secondary structure conformations. More specifically, in the poly-alanine regions, at the absence of flow, *β*-sheet and PPII dihedral conformations are on average sampled by more than 50 % of the residues and reach ~70 % at the largest flow rates we have used. Taking into account that the transition from PPII-helix to beta-sheet is very likely to happen due to their close proximity in terms of dihedrals, we can conclude that poly-alanine regions in our spidroin are primed for *β*-sheet formation by uniform flow. The predominantly proportion of alanines of ~70 % in *β*-sheet or close to *β*-sheet (that is, PPII) conformations is already very close to the 90 % percent *β*-sheet content by alanines predicted in the model of Alexandra H. Simmons et al. [63], which is based in NMR experimental studies. The occurence of torsional angles in PPII-helix conformation might result in a prefibrillar form of the repetitive region of the spidroins before forming stronger *β*-sheet interactions, as proposed by Nur Alia Oktaviani et al. [61].

### 3.2 Silk spidroin assembly under flow

Having established a coarse-grained model of the spidroin protein, MD simulations of multiple proteins offer the possibility of analyzing in detail how flow drives self-assembly. We set up systems of 25 spidroin peptides in the coarse-grained model and monitored their dynamics, interactions and oligomerization at a range of flow velocities. The simulation setup is explained in the Methods section 2.2. All-atom MD simulations were not performed in this stage due to their significantly higher computational expense and the formation of voids observed at higher velocities which limits range of accessible flow velocities [48].

As in the simulations of single chains, the silk fragment comprises ~80 residues, that is, three poly-Ala repeats and two intrinsically disorder regions. An initial pre-alignment of the peptides was performed, mimicking the multimerization at the N-terminal domains (Figure 1). The pre-alignment is maintained by tethering the peptides along the flow direction at the same position, while they can move transversely. Imposing this pre-alignment will enhance the packing of the chains (via formation of contacts mostly within the poly-Ala regions). In this way, we monitored assembly in an idealized setting with all chains in phase and subjected to uniform flow. The mean peptide fractional extension as a function of the mean flow velocity, averaged over the replicates simulated at the same flow, is shown in the Supplementary Figure S3). We covered mean fractional extensions in the range of [0.0 – 0.76]*L*. In fact, the lower four out of the total 8 velocities applied all resulted in fractional extensions close to zero. This is in sharp contrast to the single-chain simulations, in which the same range of flow velocities has lead to measurable expansions of the chain (compare Figure 2c). Here, with a relatively high density of spidroins, the lower range of flows was not sufficient to compete with the high tendency of the chains to interact and could not expand the chains prior to interchain aggregation, which further reduced their extensions drastically.

In order to analyze the self-assembly of the spidroins over time, we monitored when a certain peptide takes part of an oligomer. This was done through a clustering algorithm called Friends of Friends (FoF) (see Methods section 2.2). The criterion to form an oligomer of certain size is based on the formation of a single interchain *C_α_* – *C_α_* interaction. In Figure 4a, we visualize the oligomerization process. Data is shown for one replicate at an intermediate flow velocity, where the mean fractional extension of the peptides corresponds to 0.56L. We observe successive growth of oligomers, and an overall increase in the number of peptides involved in dimers or larger assemblies. The largest oligomer formed within the simulated time is a 10-mer in this specific case. A reduction in oligomer size is observed rarely, as fluctuations at short time scales. At a lower flow rate, with mean peptide fractional extensions close to 0.0*L*, oligomers form at shorter time scales and reach larger sizes (Supplementary Figure S4). Also, the partial reversibility observed at 0.56*L* is not observed at lower flow rates, and assembly becomes fully irreversible in this case.

**Figure 4:**
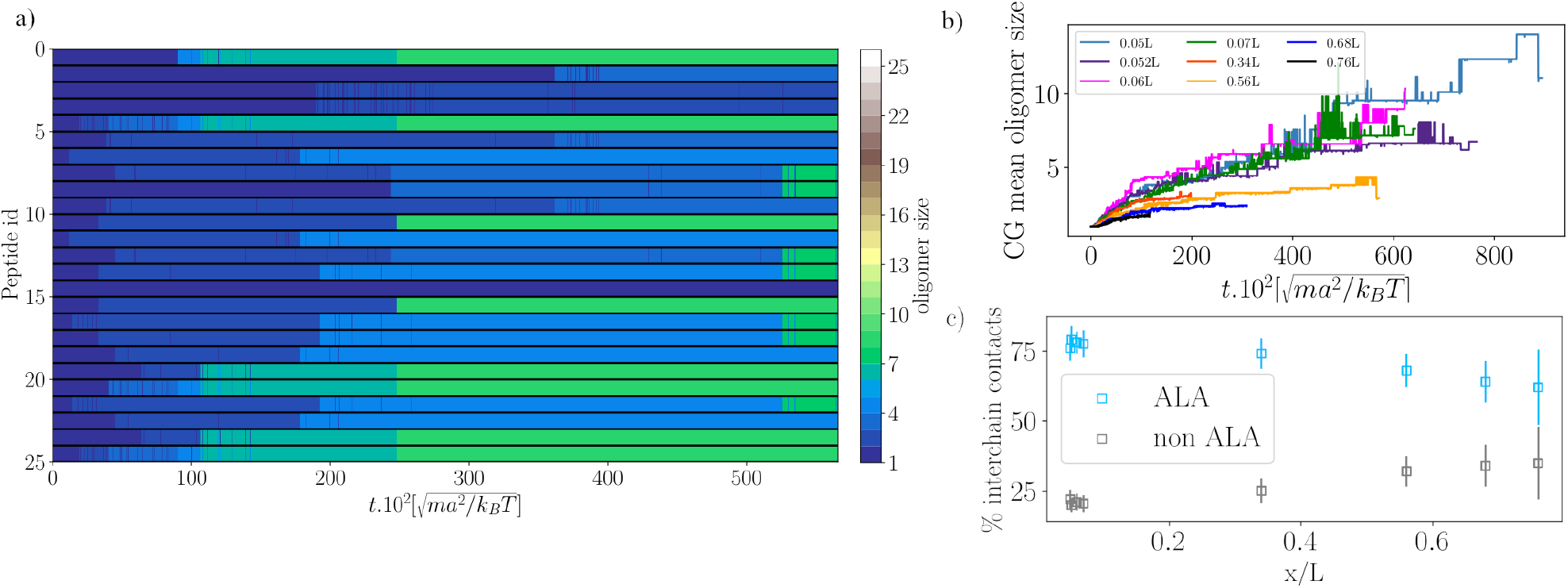
Silk oligomerization. a) Oligomer formation over time for every peptide. The color code indicates the size of the oligomer they form. Shown is the oligomerization for one replicate at a mean fractional extension of 0.56L. b) Mean oligomer size as a function of time. c) Average percentage of interchain contacts for the poly-Alanine and amorphous regions over the mean CG peptide fractional extension *x/L*.

To further quantify the effect of flow velocities on the spidroin assembly, we computed the average oligomer size as a function of time over all replicates (Figure 4b). Average oligmer sizes for individual replicates, exemplarily for two different flows, are also shown in Supplementary Figure S5). Overall, self-assembly of silk peptides slows down with increasing flow velocity (simulation snapshots for two different flows are shown in supplementary Figure S6). Slower flow causes a higher mean oligomerization at a given time (light blue, purple, green and magenta curves in Figure 4b) compared to faster flow regimes (red, orange, dark blue, and black curves in Figure 4b). This suggests that flow not only extends the spidroin chains, but in this way also reduces oligomerization. We attribute this slowdown in oligomerization to fluctuations of the protein chains orthogonal to the flow direction, which reduces the likelihood of diffusional encounters. Thus, random associations are reduced, and controlled assembly into β-sheet is enhanced. Indeed, the inter-protein contacts between the spidroins are strongly affected by flow. By definition, oligomerization involves the formation of inter-chain contacts. In addition, the type of contacts changes with flow. Namely, higher flow velocities and extensions result in more non-alanine contacts, due to the fact that chains are aligned in the oligomers (Figure 4c).

To investigate the oligomerization kinetics, we computed association and dissociation time intervals at every flow regime. Figures 5a and 5b show log-normal cumulative distributions of the time intervals at which association and dissociation events, respectively, occur. The time interval for a certain event (association or dissociation) of an oligomer of size *N_o_* corresponds to 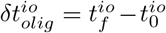. The *io* index is the oligomer identifier, 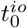 is the simulation time where an oligomer io exists and 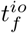 is the simulation time where either the io oligomer increased its size (association) or decreased its size (dissociation). As expected, for all flow velocities, association dominates over dissociation, reflecting the overall growth in assemblies over time (Fig. 3b). Interestingly, both the numbers of association and dissociation events increase with flow, with differences particularly large at short times. Thus, higher flow overall slows down assembly, but it does so by primarily accelerating dissociation, i.e. disassembly. Overall, flow prevents rapid aggregation into large oligomers by enhancing reversibility.

**Figure 5:**
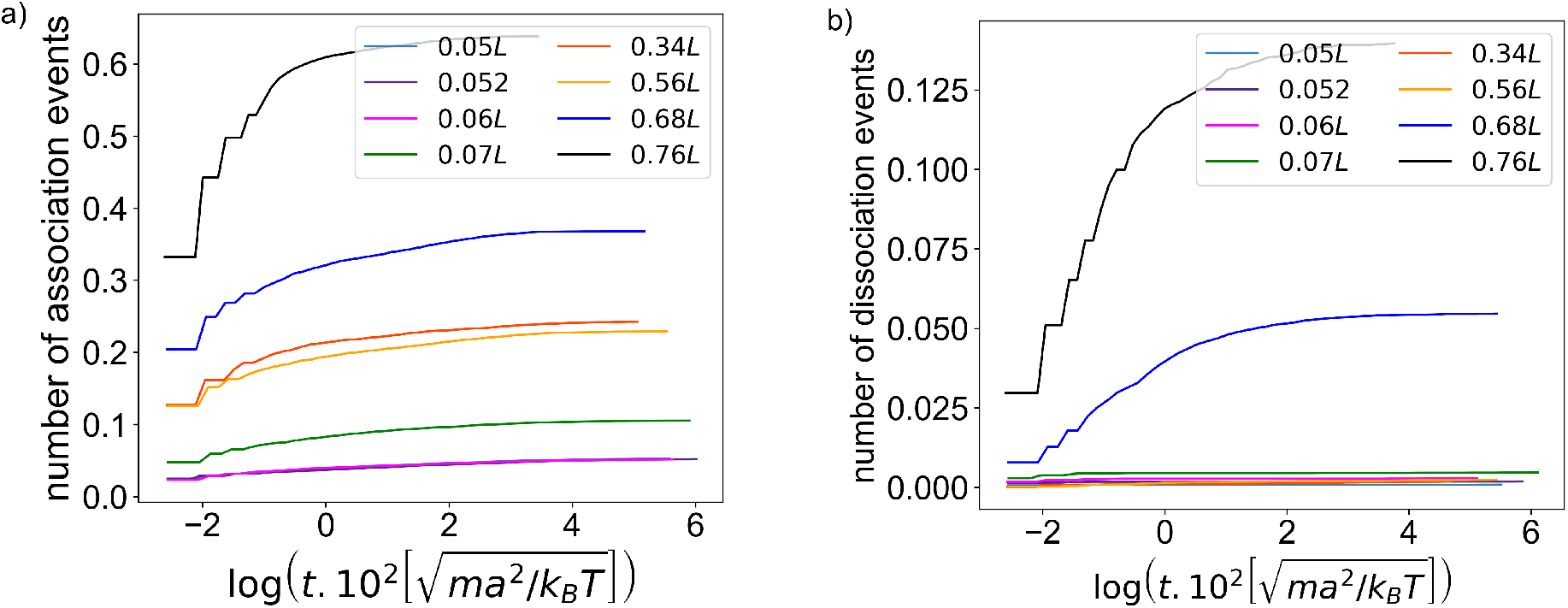
Oligomerization kinetics. Log-normal cumulative distributions of the time intervals of (a) association and (b) dissociation events, for every flow velocity (or every average protein extension). The distributions are based on data from three replicates at every flow velocity, and are normalized by the total number of time steps simulated at every flow rate. Higher flow, or larger average protein extensions, give rise to both faster association and dissociation events.

## 4 Discussion and Conclusion

We aimed to understand the influence of uniform flow on the dynamics of silk self-assembly. To this end, we focused on one-end tethered single and multiple dragline spidroins, and performed MD simulations under uniform flow on two scales: atomistic (AA) and amino-acid (CG) resolution. In this way, we could consider atomistic details from AA simulations of single chains and on the other hand covered larger time and length scales using CG simulations for the assembly. In this combination, AA simulations served to analyse in detail the structural changes of spidroins with flow (Figure 3c and Figure 3b), while the CG simulations helped us to simulate larger systems with an adequate description of the hydrodynamic interactions at play.

In the single peptide simulations, the explicit computation of the drag force from the interactions between the solvent particles and every amino acid of the spidroins allowed us to reproduce the force-extension behavior for silk proteins observed in atomic force microscopy experiments [27]. Thus, the sum of drag forces acting along a peptide by colliding water molecules can be equated with the pulling forces acting on a peptide on its termini, when estimating the resulting end-to-end distance. For the dynamic response of the spidroin protein to flow, we find a good agreement between the AA and CG model, with a loss of intrachain interactions and an increase in extension largely following the WLC model. However, the CG model is not able to reproduce the step-wise transition from a collapsed to an extended state that is observed in the AA model. In contrast, it shows a steady decrease of contacts with the mean flow velocity. We attribute this to the approximation of the spidroin as a block copolymer that lacks favorable interactions and specificity for the amorphous amino acids. We also note that the mesoscopic solvent of the CG model captures hydrodynamic interactions but the molecular nature of these interactions via hydrogen bonds and the hydrophobic effect are not explicitly included, another potential reason for the lack of a well-defined collapse-stretch transition in the CG simulations.

One interesting observation of the AA simulations is the increase of alanines in beta-sheet conformations with flow, increasing their propensity for fibrillation and the formation of beta-sheet crystals as found in silk fibers [55].

Due to the high computational cost and the limited flow velocity that can be achieved in AA MD simulations [48], we only simulated CG multiple spidroins to monitor their assembly. Silk fiber formation is a highly complex and multiscale process occuring in the gland, and capturing the full complexity, from the gradual increase in the flow rate, the transition from a micellar to an extended state of the proteins, to the change in pH and salt concentration, to only name a few, is prohibitive. Our highly simplified setup was motivated by the following scenario: The N and C-terminal domains (NT and CT) experience conformational changes upon assembly inside the S-gland. The CT is dimerized from the tail of the gland [21], and the NT domain dimerizes inside the first part of the s-duct after a drop of pH [64]. Therefore, both terminal domains promote the initial interconnections of the proteins, and thereby their pre-alignment. To mimic this role of the terminal domains of the spidroins for multimerization (Figure 1), we pre-aligned the repetitive silk peptides to maintain them in phase. This setup facilitated the assembly via crystal formation in the poly-Ala regions, and allowed to focus mostly in the flow effects involved in the spidroins’ packing.

Regarding the assembly by the formation of crystals, several studies have pointed out the high tendency of *β*–sheet structures between the poly-Ala regions of the spidroins [63, 65, 66]. CG simulations can extend the limited time scales of the AA systems, but at the expense of an accurate structural model. We here chose a CG model that still captures the sidechain packing of alanines across beta-sheets in crystals by using two beads per amino acid. Our CG model was able to identify two key tendencies of flow-induced silk fiber assembly. Flow extends the chains and overall slows down the oligomerization. The underlying kinetics shows that this is a consequence primarily of enhancing and accelerating also dissociation. As a result, flow introduces reversibility and removes kinetic traps. This situation is similar to assembly of protein complexes under quiescent conditions, like virus capsids, which is stalled in kinetic traps if the free energy gain is too high [67, 68, 69]. For the case of silk discussed here, moderate amounts of shear, but not too high values increase the quality of the assembly, allowing proper beta-sheet configurations to form not only because of the mere extension and alignment of the poly-alanine repeats, but also by speeding up their reversible association and dissociation, as a mechanism of quality control. We note that our observations on silk assembly at the CG level, by comparison to the all-atom simulations, correspond to flows in the range of 0.5-3 m/s (Figure 2c). These speeds are relevant speeds, albeit at the upper boundary, of those used in a laboratory setting or observed within in the spider gland [70]

Overall, we have investigated by computer simulations the effect of flows on spider silk peptides. Our study gives insights into their flow-induced non-equilibrium conformational dynamics and how flow influences their self-assembly. Microfluidic experiments have shown how elongational flow enhances *β*–sheet formation [18], which we here can confirm. However, due to the high hydrophobicity of the silk dope, a careful control of the solvent during the assembly process is required. Microscopy experiments under controlled conditions of pH and flow to prevent fatal aggregation, and where the assembly of the spidroins can be monitored in real-time are much desired. Such experiments can help to put the mechanisms of flow-induced *β*–sheet formation we here put forward to test.

## 5 Acknowledgments

This work was supported by the Klaus Tschira Foundation. We also acknowledge funding through the Deutsche Forschungsgemeinschaft (DFG, German Research Foundation) under Germany’s Excellence Strategy – 2082/1 – 390761711 (cluster of excellence 3DMM2O) and through research grant GR 3494/7-3. This research was conducted within the Max Planck School Matter to Life supported by the German Federal Ministry of Education and Research (BMBF) in collaboration with the Max Planck Society. Computing time was allocated by the state of Baden-Wörttemberg through bwHPC and DFG through grant INST 35/1134-1 FUGG.

## Notes

### Competing Interest Statement

The authors have declared no competing interest.

