## supplemental figures for "The role of hydrodynamic flow in the self-assembly of dragline spider silk proteins"

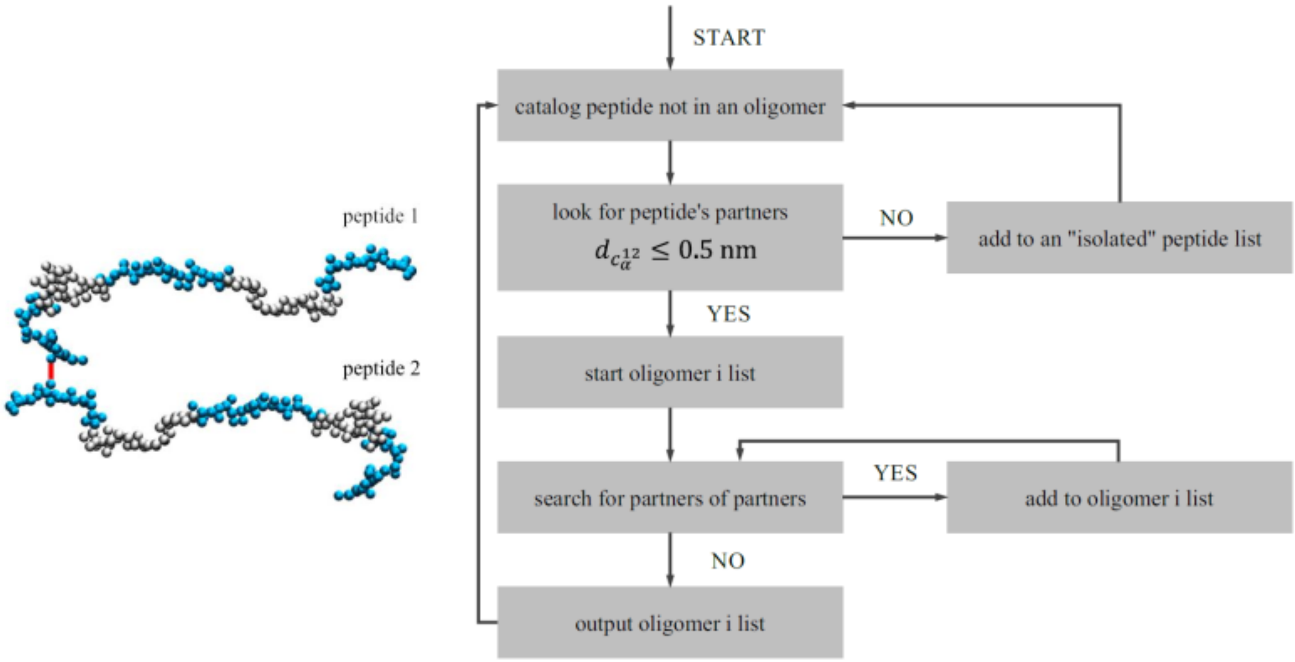


**Figure S1:** Scheme of the Friends of Friends algorithm

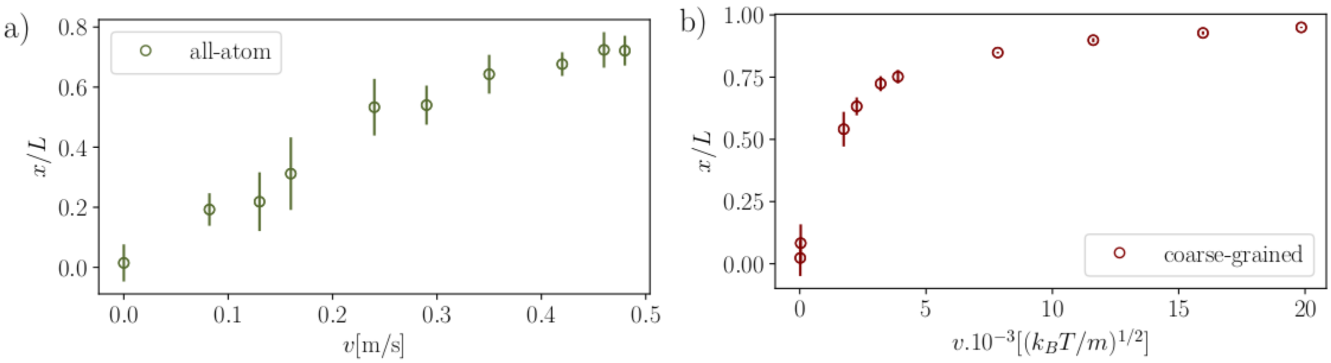


**Figure S2:** Fractional extension over flow rate for all-atom (a) and coarse-grained (b) simulations. As expected, for both the atomistic and coarse-grained scales, we obtain on average larger extensions at larger flow velocities. The flow velocities used in the coarse-grained simulations cover a wider range and reach the regime of largely extended peptide conformations ($x/L \sim 0.9$) already from medium flow velocities onwards.

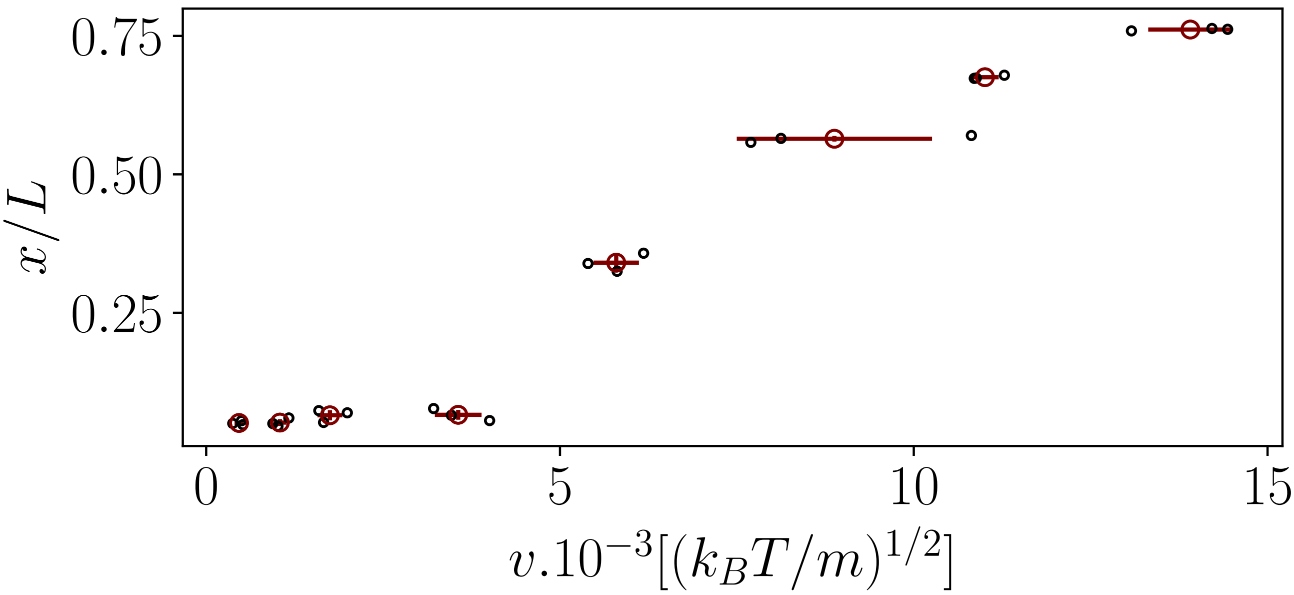


**Figure S3:** Mean CG peptide fractional extension $x/L$ as a function of the mean flow velocity. Black points correspond to individual replicates, and maroon points correspond to the average over replicates in every flow regime, considering the last half of every trajectory.

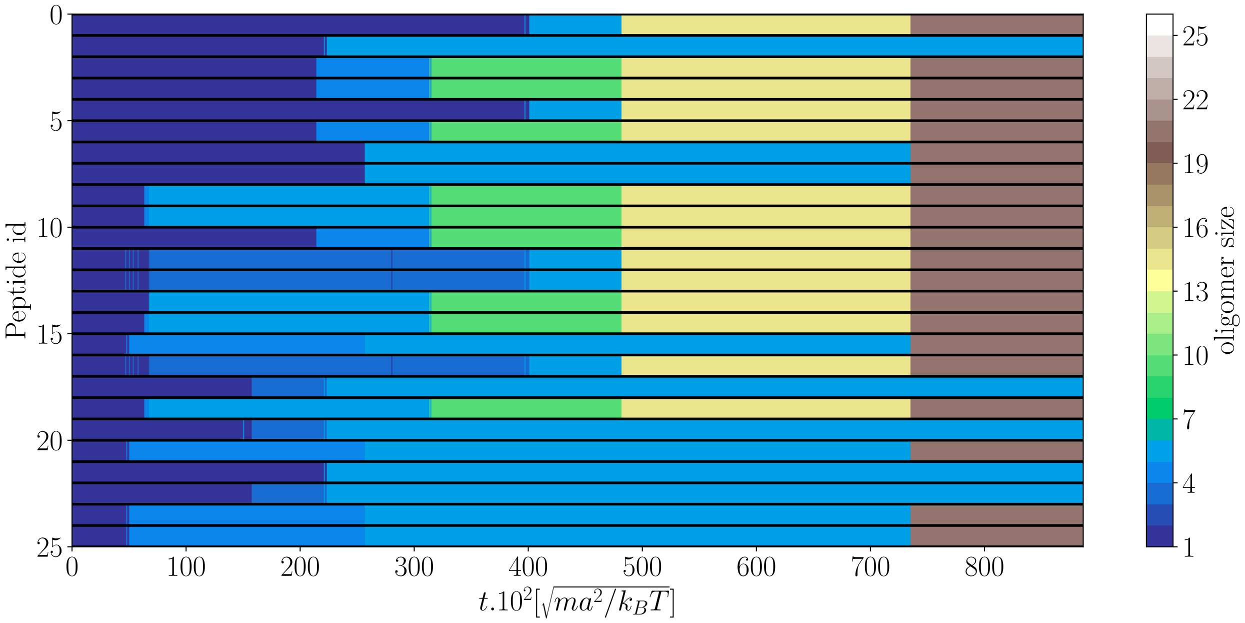


**Figure S4:** Oligomer formation in time for every peptide, depending on the size of the oligomer they form. The figure is shown for one replicate at a mean fractional extension of 0.05L.

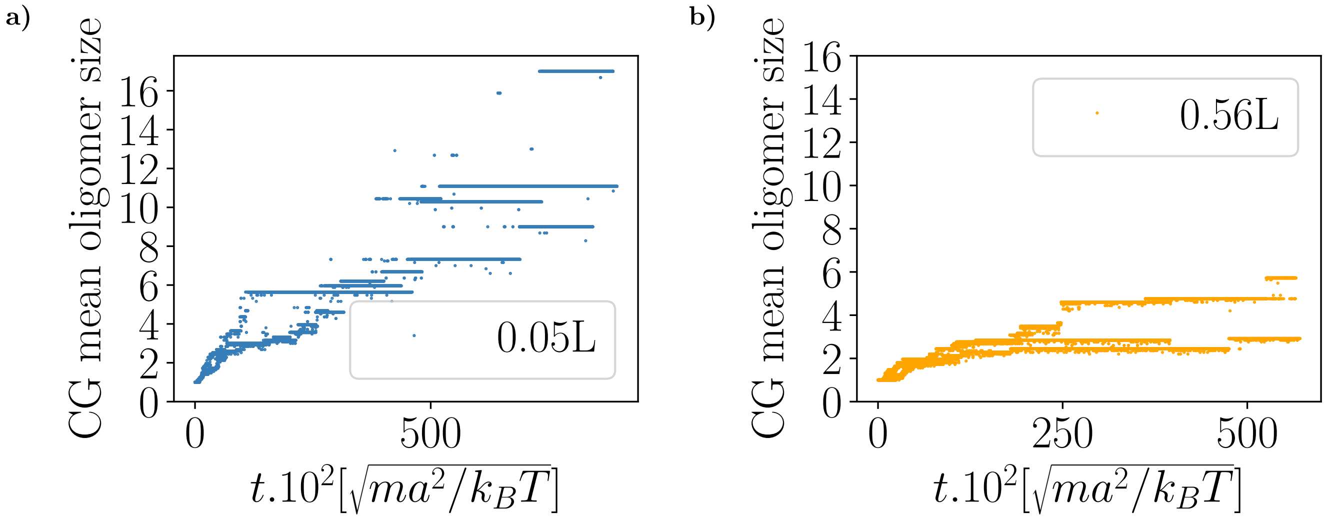


**Figure S5:** Mean oligomer size per replicate as a function of time for every mean fractional extension

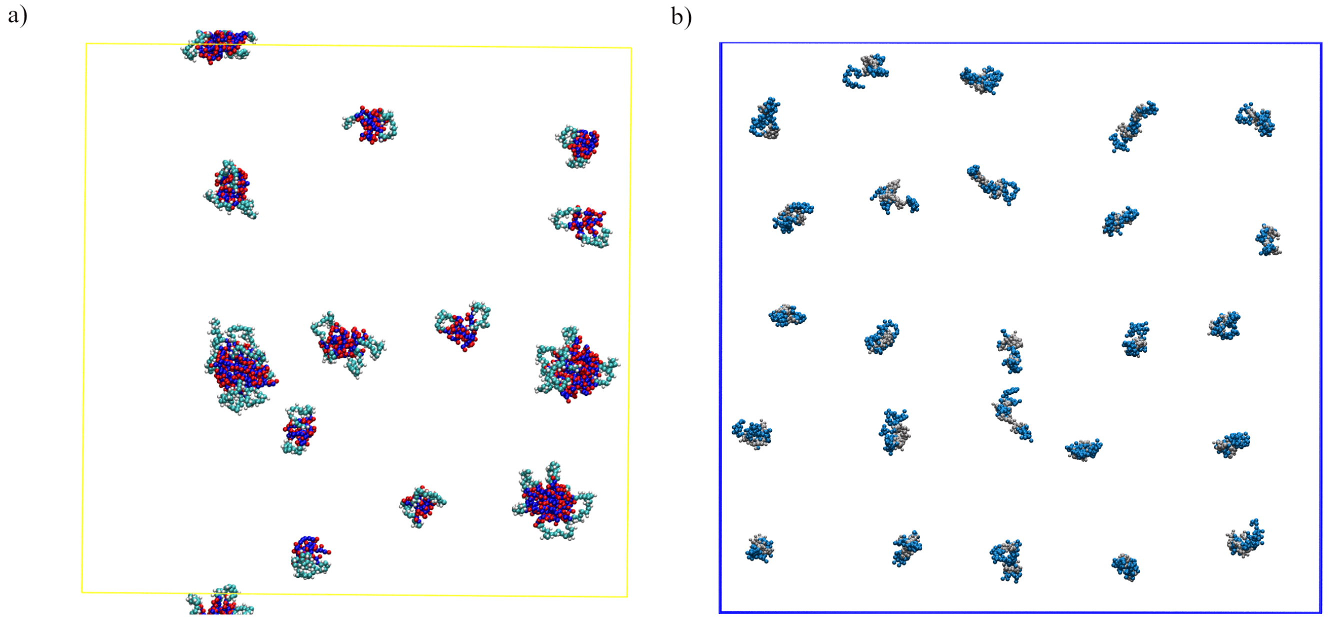


**Figure S6:** Transversal view of spidroins' assembly at (a) the lowest flow rate, corresponding to an initial mean fractional extension of $0.05L$ and (b) the highest flow rate, corresponding to an initial mean fractional extension of $0.76L$. The snapshots of the simulations were taken at a MPCD time of 6.25x10$^2$ $\left[\sqrt{ma^2/k_BT}\right]$.
